# Effect of the S2’ site cleavage on SARS-CoV-2 spike

**DOI:** 10.1101/2025.04.02.646838

**Authors:** Wei Shi, GM Jonaid, Md Golam Kibria, Jacob Allen, Hanqin Peng, Sophia Rits-Volloch, Haisun Zhu, Shaowei Wang, Richard M. Walsh, Jianming Lu, Bing Chen

## Abstract

SARS-CoV-2 initiates infection of host cells by fusing its envelope lipid bilayer with the cell membrane. To overcome kinetic barriers for membrane fusion, the virus-encoded spike (S) protein refolds from a metastable prefusion state to a lower energy, stable postfusion conformation. The protein is first split into S1 and S2 fragments at a proteolytic site after synthesis, and presumably further cleaved at a second site, known as the S2’ site, before membrane fusion can occur. We report here a cryo-EM structure of S2 fragment after the S2’ cleavage, possibly representing a late fusion intermediate conformation, in which the fusion peptide and transmembrane segment have yet to pack together, distinct from the final, postfusion state. Functional assays demonstrate that the S2’ cleavage accelerates membrane fusion, probably by stabilizing fusion intermediates. These results advance our understanding of SARS-CoV-2 entry and may guide intervention strategies against pathogenetic coronaviruses.

## Introduction

SARS-CoV-2, an enveloped RNA virus, enters a host cell by fusing its envelope membrane with the target cell membrane. Although energetically favorable, the membrane fusion reaction requires free energy to overcome high kinetic barriers caused primarily by the repulsive hydration force when the two membranes approach each other ^1,2^. The catalyst of the fusion reaction is the virus-encoded spike (S) protein that refolds from a high-energy, metastable prefusion conformation to a low-energy, stable postfusion state during the fusion process^3–5^.

The S protein is a heavily glycosylated type I membrane protein anchored in the viral membrane. It is a class I viral fusion protein, like HIV-1 envelope glycoprotein, influenza hemagglutinin or Ebola glycoprotein^3,6^. The S protein is first synthesized as a single-chain precursor that trimerizes and subsequently undergoes cleavage at a polybasic proteolytic site into two fragments: S1 for receptor-binding and S2 for membrane fusion^6–8^. This proteolytic site was introduced when SARS-CoV-2 acquired a four-residue insertion (Pro-Arg-Arg-Ala) at the S1/S2 boundary for recognition by the host protease furin in the trans-Golgi network of the virus-producing cells^8,9^. SARS-CoV lacks this site^10^. The furin cleavage primes the S protein, making the prefusion structure metastable with respect to the postfusion conformation. The cleavage appears to promote infection of lung cells by SARS-CoV-2; it also contributes to viral pathogenesis in animal models^8,11–13^. To infect a new host cell, the furin-cleaved S trimer binds to the entry receptor ACE2 (angiotensin converting enzyme 2) on the cell surface. It is believed to be further processed at a second proteolytic site in S2, designated S2’ site, by another host protease – either TMPRSS2 (transmembrane serine protease 2) at the cell surfaces, or cathepsin L in endosomes after endocytosis^9,14,15^. The S2’ site cleavage is generally thought to be the key trigger for activating the S trimer to undergo large conformational changes leading to membrane fusion. TMPRSS2, a member of the TMPRSS family of proteases, is a type II transmembrane serine protease that cleaves at an arginine or lysine residue^16,17^. It is present in many epithelial tissues, but only nasal, bronchial, and gastrointestinal epithelia express both ACE2 and TMPRSS2^18–20^. Cathepsin L, a cysteine protease responsible for protein degradation in the late endosomes and lysosomes, shows little sequence specificity for its cleavage sites^21^. It is expressed in all tissues and cell types, and associated with chronic inflammation^22–24^. Proteolytic activation of spike, at least by TMPRSS2, as well as downstream membrane fusion seem to require both ACE2 engagement and pre-cleavage at the S1/S2 junction^9^.

Once activated, the prefusion S trimer undergoes large and irreversible rearrangements to fold into the stable postfusion structure, inducing membrane fusion^15^. Previous structural studies have revealed many molecular details of both the prefusion and postfusion conformations of the SARS-CoV-2 spike^25–30^. The prefusion S trimer samples either a closed and symmetrical structure with three receptor binding domains (RBDs) in the down conformation representing a receptor-inaccessible state, or an open and asymmetrical structure with one RBD in the up conformation that represents a receptor-accessible state^25,26^. In the postfusion state, S1 dissociates as a monomer and the S2 trimer adopts a rigid, rod-like shape. The membrane-interacting regions, functionally critical for membrane fusion but missing in all the previous structures, are visible in a cryo-EM structure of the intact postfusion S2 converted from the prefusion S trimer in lipid nanodiscs after binding to ACE2^30^. There are nine membrane-spanning helices (three per protomer) in the lipid bilayer region, including the fusion peptide (FP), which forms a hairpin-like wedge of two membrane-spanning helices, and the transmembrane (TM) helix, which wraps around the FP. Comparison of the prefusion and postfusion structures indicates that formation of the long postfusion central helix translocates the FP by ∼185 Å to insert into the target membrane^30^. Further refolding events to form the stable postfusion bundle effectively bring the viral and target cell membranes in close apposition and allow the interactions between the FP and TM to complete membrane fusion. Nonetheless, the cryo-EM postfusion structure does not explain the functional role of the S2’ site cleavage, as the site is disordered in the structure and as biochemical evidence indicates it is uncleaved^30^, suggesting that formation of the S2 postfusion conformation does not require cleavage at the S2’ site.

To assess the effect of the S2’ cleavage on the structure and function of SARS-CoV-2 spike protein, we have prepared an S2 fragment with the S2’ site cleaved (referred to as “S2’ fragment” thereafter) and determined its structure by cryogenic electron microscopy (cryo-EM). The S2’ fragment tends to form clusters of 2-3 trimers through interactions between their hydrophobic FPs. The S2’ cleavage does not alter the location of the FP, but the segment immediately downstream of the S2’ site (S^816–834^), which was widely accepted previously as the fusion peptide^31–33^, interacts with the membrane proximal external region of spike protein to help stabilize an asymmetrical postfusion-like structure. Functional assays demonstrate that S2’ cleavage by TMPRSS2 is not essential for membrane fusion, but significantly accelerates the membrane fusion reaction.

## Results

### Production of an S2 fragment with the S2’ site cleaved

To assess the structural impact of the S2’ cleavage, we developed a protocol for producing an S2 fragment with the S2’ site fully cleaved (S2’ fragment; Fig. 1A) by trypsin, which is more effective and specific than soluble TMPRSS2^34^. We confirmed that addition of trypsin in a cell-cell fusion assay enhanced fusion between S-expressing and ACE2-expressing cells, equivalent to co-expressing the membrane-bound TMPRSS2 with ACE2 in the target cells (Fig. S1A), suggesting that trypsin can indeed functionally replace TMPRSS2, as reported for other coronavirus spikes^35,36^. To prepare the S2’ fragment, we first incubated the HEK293T cells stably expressing the C-terminally strep-tagged, full-length SARS-CoV-2 spike protein with soluble ACE2 and then treated the cells with trypsin, which could not cleave the spike protein in the absence of ACE2 under the same conditions (Fig. S1B). When we extracted the S protein by detergent dodecyl-β-D-maltoside (DDM) and purified it using a strep-tactin column, an intact S2’ fragment, as confirmed by N-terminal sequencing and the presence of C-terminal strep tag from western blot, was obtained (Fig. 1B-1D). It migrated as a band of ∼75 kDa by SDS-PAGE, smaller than both S1 and S2 fragments, as expected (Fig. S1C). Negative stain EM analysis showed that the purified S2’ fragment had adopted a postfusion-like conformation with the characteristic rod-like shape, either as individual trimers or as clusters of 2 or 3 trimers (Fig. 1E). These S2’-cleaved trimers resembled in both shape and length the postfusion trimer formed by the uncleaved S2 (Fig. S1D; ref^30^). Clusters of trimers were also found with the uncleaved S2 trimer in detergent or the S2’ trimer reconstituted in nanodiscs (Fig. S1D and S1E).

**Figure 1.**
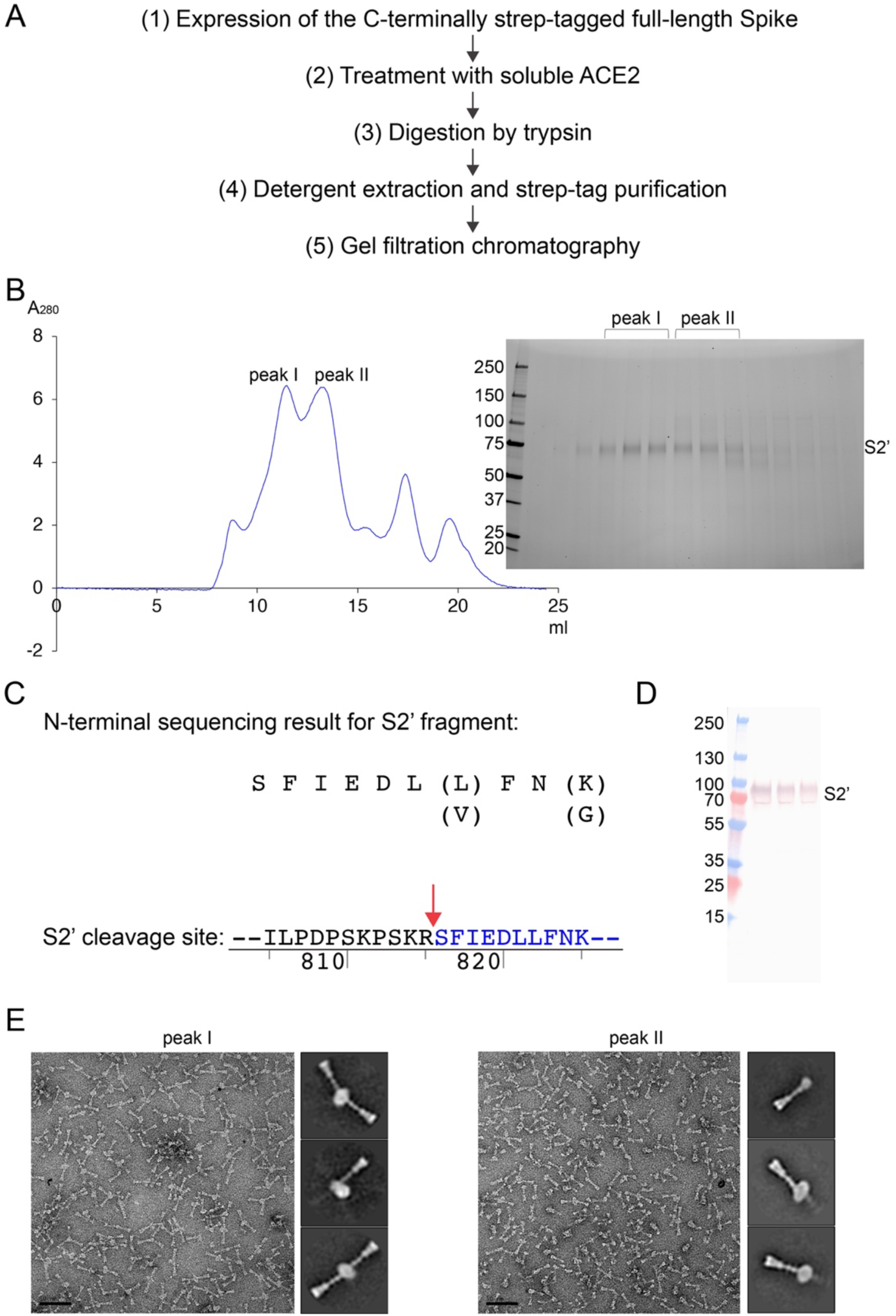
Preparation of the S2’ fragment of SARS-CoV-2 spike protein. (**A**) Strategy to purify an S2 fragment with the S2’ site cleaved – S2’ fragment. (**B**) The purified intact SARS-CoV-2 S2’ fragment in DDM was resolved by gel-filtration chromatography on a Superose 6 column. Two major protein peaks contain a fragment of ∼75 kDa, as analyzed by SDS-PAGE. (**C**) N-terminal sequencing for the S2’ fragment conforming the correct cleavage. The sequence of the S2’ cleavage site is also shown with the cleavage position indicated by a red arrow. (**D**) Western blot using anti-strep tag antibody confirming the presence of the C-terminal strep tag. (**E**) Negative stain EM of the two major peaks: Peak I – mainly clusters of postfusion spike trimers and Peak II - mainly individual postfusion spike trimers. Representative image and 2D averages are shown, and the box size of 2D averages is ∼460Å.

### Cryo-EM structure of the S2’ fragment

We next determined by cryo-EM the structure of the S2’ trimer solubilized in DDM. We recorded cryo-EM images on a Titan Krios electron microscope equipped with a Falcon 4i direct electron detector and used cryoSPARC^37^ for particle picking, two-dimensional (2D) classification, three-dimensional (3D) classification and refinement (Fig. S2). We obtained one major class from 3D classification and refined it to 3.1Å resolution (Fig. S2-S4; Table S1). To further improve the local resolution near the transmembrane region embedded in detergent micelles, we performed additional masked local classification and refinement, leading to a 3.0 Å map covering the fusion peptides and TM segments.

The core structure of the S2’ trimer in the postfusion-like conformation, formed by the helices of HR1, HR2, CH and 3H (Fig. 2A and 2B), is nearly identical to those of the previous postfusion spike structures^27,30,38^. The S1/S2-S2’ fragment, which includes the 3H segment and is no longer covalently linked to the rest of S2 after the S2’ site has been cleaved (Fig. 1A), remains tightly associated with the trimer. HR1, CH and part of the FP form a long central coiled-coil that is reinforced by helix-bundles with HR2 and 3H. Another segment of the S1/S2-S2’ fragment, β^718–729^, forms a three-stranded β sheet with the connector domain (CD) that wraps around the C-terminal end of the CH coiled-coil. The C-terminal half of S2’ folds back onto the central coiled-coil, with the strand β^1127–1135^ joining the connector β sheet, the short helix α^1148-1155^ packing between two neighboring 3Hs, and the HR2 helix bundling with the HR1 coiled-coil, to effectively bring the TM segment and FP at the same end of the trimer (Fig. 2C).

**Figure 2.**
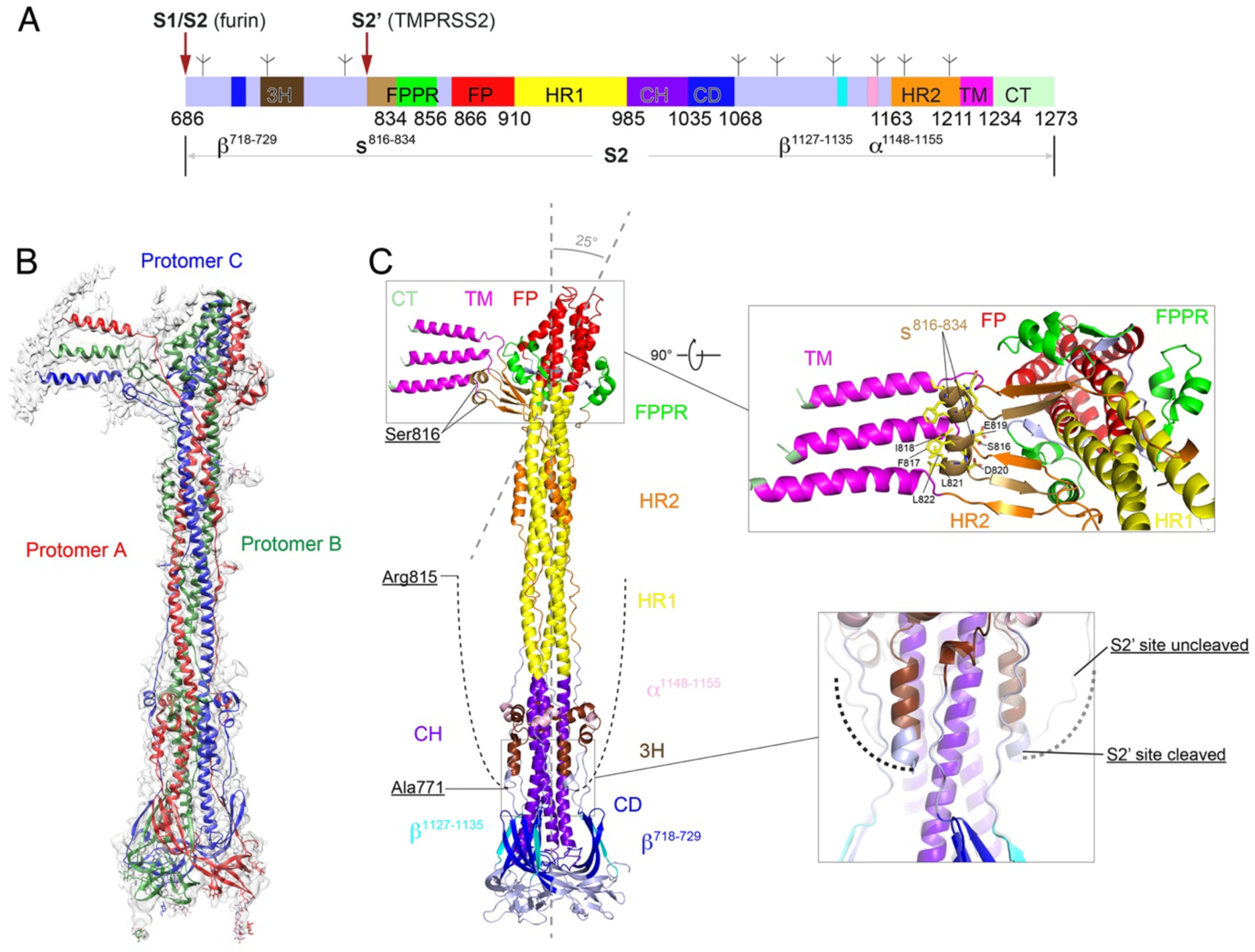
Structure of the S2’ fragment of SARS-CoV-2 spike in the postfusion conformation. (A) A schematic of the SARS-CoV-2 S2 fragment, which has the following segments: S1/S2, the furin cleavage site at the S1/S2 boundary; S2’, S2’ cleavage site; β^718-729^, a β-strand formed by residues 718-729 in the S1/S2-S2’ fragment; 3H, three-helix segment; s^816-834^, a segment containing residues 816-834; FPPR, fusion peptide proximal region; FP, the fusion peptide; HR1, heptad repeat 1; CH, central helix region; CD, connector domain; β^1127-1135^, a β-strand formed by residues 1127-1135; α^1148-1155^, an α-helix formed by residues 1148-1155; HR2, heptad repeat 2; TM, transmembrane anchor; CT, cytoplasmic tail; and tree-like symbols for glycans. Also see ref^30^. (B) The structure of the S2’ trimer fits into a 3.1Å density map. Three protomers (A, B, C) are colored in green, blue and red, respectively. (**C**) Overall structure of the S2’ trimer in the postfusion conformation shown in ribbon diagram. Various structural components in the color scheme shown in **B** include β^718-729^, a β-strand formed by residues 718-729 in the S1/S2-S2’ fragment; 3H, three-helix segment; s^816-834^, a segment containing residues 816-834; FPPR, fusion peptide proximal region; FP, the fusion peptide; HR1, heptad repeat 1; CH, central helix region; CD, connector domain; β^1127-1135^, a β-strand formed by residues 1127-1135; α^1148-1155^, an α-helix formed by residues 1148-1155; HR2, heptad repeat 2; TM, transmembrane anchor; and CT, cytoplasmic tail. The segment immediately upstream of the S2’ cleavage site (residues 772-815) is disordered and indicated by a dashed line. A close-up view of the transmembrane region after a 90° rotation is also shown and the residues of the structured s^816-834^ segment are indicated. Superposition of the C-terminal region of 3H from the S2’ trimer (colored) and the uncleaved S2 trimer (gray) is shown with an extra helical turn packing against the CH coiled-coil in the S2’ trimer.

The structure deviates most markedly from the postfusion S2 structure at the end that includes the membrane interacting segments – the fusion peptides and the transmembrane segments (Fig. 2C). The local threefold axis of the fusion peptide cluster and adjacent structures, including the FPPR, tilts by about 25° with respect to the threefold axis of the rest of the trimer. Moreover, the three TM helices and the parts of the HR2 segments N-terminal to them project laterally, rather than surrounding the cone-like FP cluster, and S^816-834^ segments from two of the three subunits reinforce the lateral projection. The N-terminal part of this segment forms an amphipathic helix on the surface of the detergent micelle surrounding the hydrophobic TM helices, which run roughly parallel to each other and may also interact as the distances of some of their hydrophobic side chains are within the range (3-4Å) for hydrophobic interactions (Fig. S5). The two fully-structured S^816-834^ segments also form a complex set of β-strand interactions with the TM-proximal HR2 regions: one forms a three-stranded β sheet with the two HR2 strands and the other interacts with the third connecting HR2 strand (Fig. 2C). This asymmetrical structure may represent a conformation preceding the final symmetrical postfusion conformation in our previous cryo-EM structure, in which the TM helices wrap symmetrically around the FP^30^. The two amphipathic helices of the S^816-834^ segments would then dip into the viral membrane immediately before its fusion with the target cell membrane, into which the FP has inserted. Toward the opposite end of the rod, the S2’ trimer has gained, with respect to the uncleaved S2 trimer, one extra turn of helix near Ile770 at the end of 3H, packing against the CH coiled-coil (Fig. 2C). This local change is probably due to the relaxed structural constraints after the cleavage that allows such folding. Overall, the new structure shows that the FP location is not affected by the S2’ site cleavage and that the S1/S2-S2’ fragment does not dissociate from the trimer after the cleavage.

We have also obtained a low-resolution map from the same data set and from an independent data set for a dimer of S2’ trimers formed by interactions between their FP regions (Fig. 3). This interaction may also contribute the tilting of the three-fold axis of the FPs of the interacting trimers (Fig. 2C). The density for one trimer is substantially stronger than that of the other trimer, confirming the heterogeneity at the interface that may explain the accommodation of a third trimer as shown by negative stain EM (Fig. 1E).

**Figure 3.**
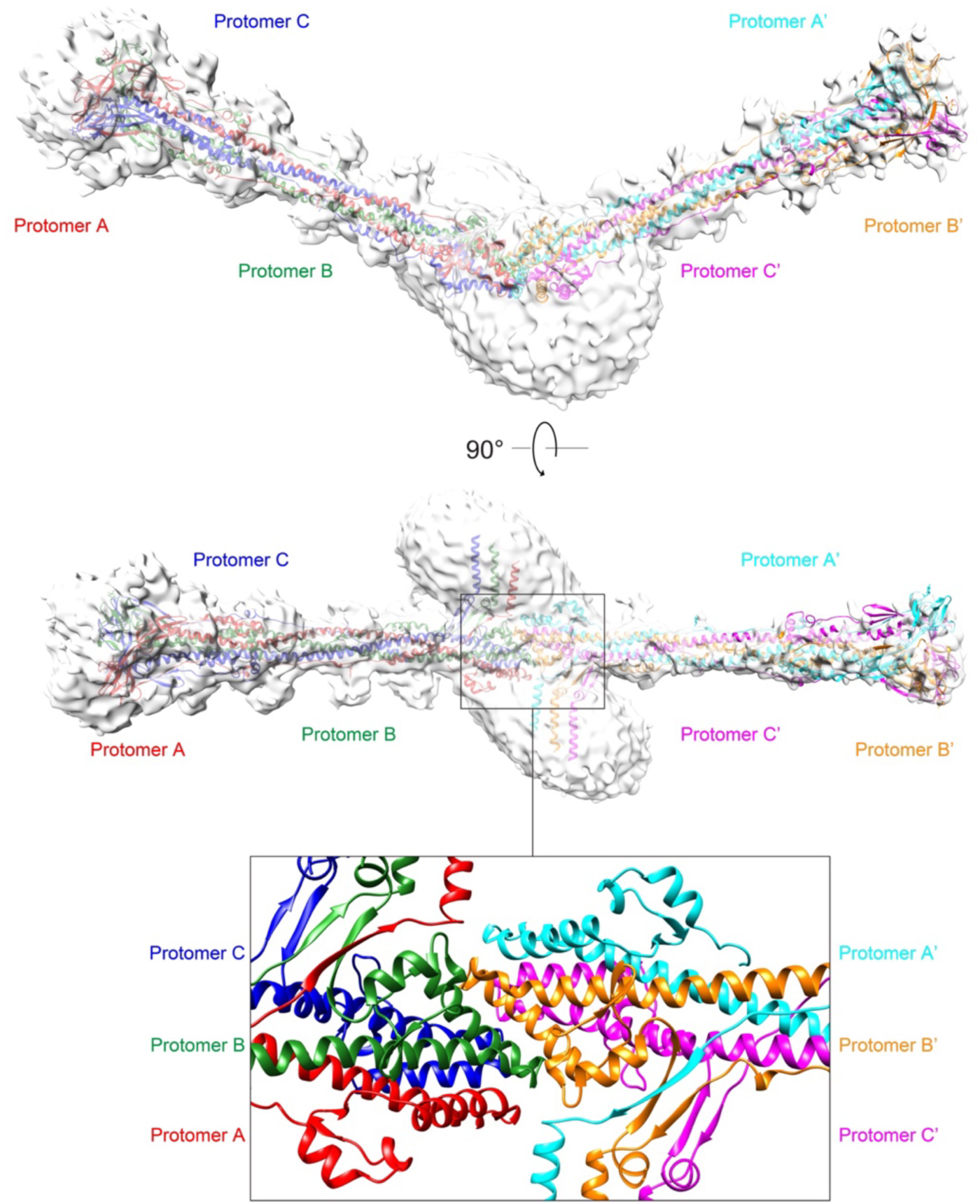
Cryo-EM map and model of a dimer of the S2’ trimer. Top and middle, a model of the dimer of S2’ trimer fits into a 6.8Å density map. Three protomers (A, B, C) for one trimer are colored in red, green and blue, respectively; and three protomers (A’, B’, C’) for another trimer are colored in orange, magenta and cyan, respectively. Bottom, a close-up view of the FP regions from the two trimers that mediate dimerization.

### R815 in the S2’ cleavage site is dispensable for membrane fusion

The S2’ cleavage site was identified in SARS-CoV spike by mutagenesis, but the impact of the mutations on the stability of the prefusion trimer was not fully addressed^39^. As shown in Fig. S6, Arg815 at the S2’ site in the prefusion SARS-CoV-2 spike trimer probably forms a salt bridge with Asp867. There may be a hydrogen bond between Arg815 and Ser813, and also a transient cation-ρε interaction between Arg815 and Phe823, making Arg815 a critical residue for stabilizing the local structure of the prefusion trimer. We mutated Arg815 in the SARS-CoV-2 spike to all other possible residues and found that many substitutions, which did not alter the S expression but would block the S2’ site cleavage by TMPRSS2 or trypsin, still efficiently supported cell-cell membrane fusion (Fig. 4A and Fig. S7A). The mutations that completely abolished membrane fusion also disrupted the prefusion trimer structure, as demonstrated by negative stain EM images of the purified mutant proteins (Fig. S8A) and by major changes in their antigenic properties (Fig. S8B). As a control, we also mutated the furin cleavage site at the S1/S2 boundary; the mutant (FSmut) had ∼60-70% of the fusion activity of the wildtype spike, consistent with the previously published results^11,40,41^. These data demonstrate that Arg815 helps stabilize the prefusion conformation, but the specific cleavage at the S2’ site involving this residue is not essential for membrane fusion.

**Figure 4.**
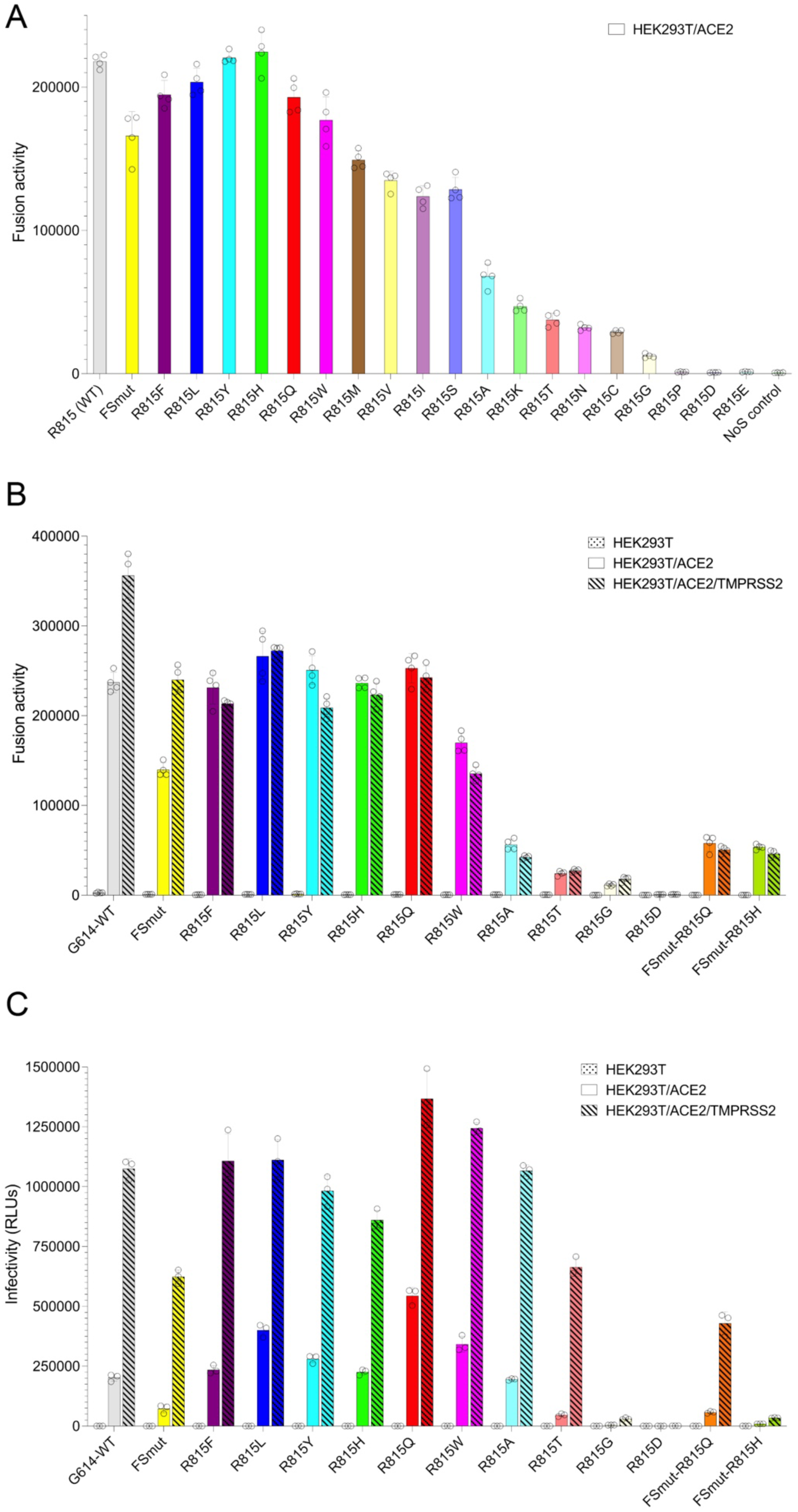
Effect of mutations in the S2’ cleavage site. (**A**) HEK293T cells transfected with the full-length S protein (G614) and its mutants were tested for membrane fusion with the ACE2-expressing cells in a β-galactosidase-based cell-cell fusion assay. FSmut, furin site mutant; NoS, a negative control without S protein. The experiment was repeated at least three times with similar results. Error bars, mean ± s.d. (**B**) HEK293T cells transfected with the full-length S protein (G614) and selected mutants were tested for membrane fusion with HEK293T cells (negative control), or the ACE2-expressing cells, or the ACE2-and TMPRSS2-expressing cells in the β-galactosidase-based cell-cell fusion assay. FSmut, furin site mutant; FSmut-R815Q and FSmut-R815H are the mutants with both the furin site and S2’ site mutated. The experiment was repeated at least three times with similar results. Error bars, mean ± s.d. (**C**) Infection of HEK293T cells (negative control), or cells transfected with ACE2 or with both ACE2 and TMPRSS2 by SARS-CoV-2 virus like particles (VLPs) produced using the full-length G614 or selected mutants, as indicated. The experiment was repeated three times with similar results. Error bars, mean ± s.d.

HEK293T cells express little endogenous TMPRSS2; overexpression of the enzyme substantially enhanced cell-cell fusion by the wildtype spike and also the FSmut, as expected (Fig. 4B). The Mutating the residue Arg815 completely eliminated TMPRSS2-mediated enhancement, whether or not the furin cleavage site had been mutated (Fig. 4B), suggesting that the S2’ site cleavage by TMPRSS2 was necessary for enhancing fusion. When these mutated constructs were tested in a SARS-CoV-2 virus-like particle (VLP) system^42^, VLP infectivity was still enhanced by TMPRSS2 (Fig. 4C). Several S mutants (R815F, R815L, R815H, R815Q and R815W) efficiently incorporated into the VLPs and showed infectivity comparable to that of the wildtype spike (Fig. 4C and Fig. S7B). The two mutations (R815P and R815D) that disrupted the prefusion structure of S trimer also prevented S incorporation into the VLPs and led to no infectivity. All other fusion-active mutants showed TMPRSS2-dependent enhancement in the VLP infectivity regardless of whether the S2’ site or furin site or both had been mutated or not (Fig. 4C). TMPRSS2-mediated internalization of both SARS-CoV and SARS-CoV-2 has been reported^43,44^, but the underlying mechanism remains unknown. Our current data show that TMPRSS2-mediated enhancement in VLP infectivity, probably via the endosomal entry pathway, does not depend on cleavage at the S2’ site, unlike membrane fusion at the cell surface.

### S2’ cleavage by TMPRSS2 accelerates S-mediated membrane fusion

To further assess the impact of the S2’ site cleavage by TMPRSS2, we analyzed the fusion activities of various S constructs in the presence of the TMPRSS2 inhibitor, camostat, and the cathepsin L inhibitor, E64-d. In the HEK293T cells expressing ACE2 without TMPRSS2, the fusion activity of the wildtype spike was not affected by either camostat or E64-d or both combined (Fig. 5A). The same pattern was observed for the FSmut, R815Q and FSmut-R815Q, although with lower fusion activities for the constructs with the furin site mutated. In the HEK293T cells expressing both ACE2 and TMPRSS2, the fusion activity of the wildtype spike was enhanced, as expected, but the enhancement could be completely suppressed by camostat alone (Fig. 5A). FSmut showed a similar pattern but with an overall lower fusion activity. The two constructs with the S2’ site mutated, R815Q and FSmut-R815Q, were insensitive to all the inhibitors. These results indicate that there is no detectable endogenous TMPRSS2 activity in HEK293T cells that could enhance S-mediated membrane fusion, and that other host protease(s) but not cathepsin L may be responsible for the fusion activity detected in the ACE2 only-expressing cells if a proteolytic event near the S2’ site is required for activating the spike protein.

**Figure 5.**
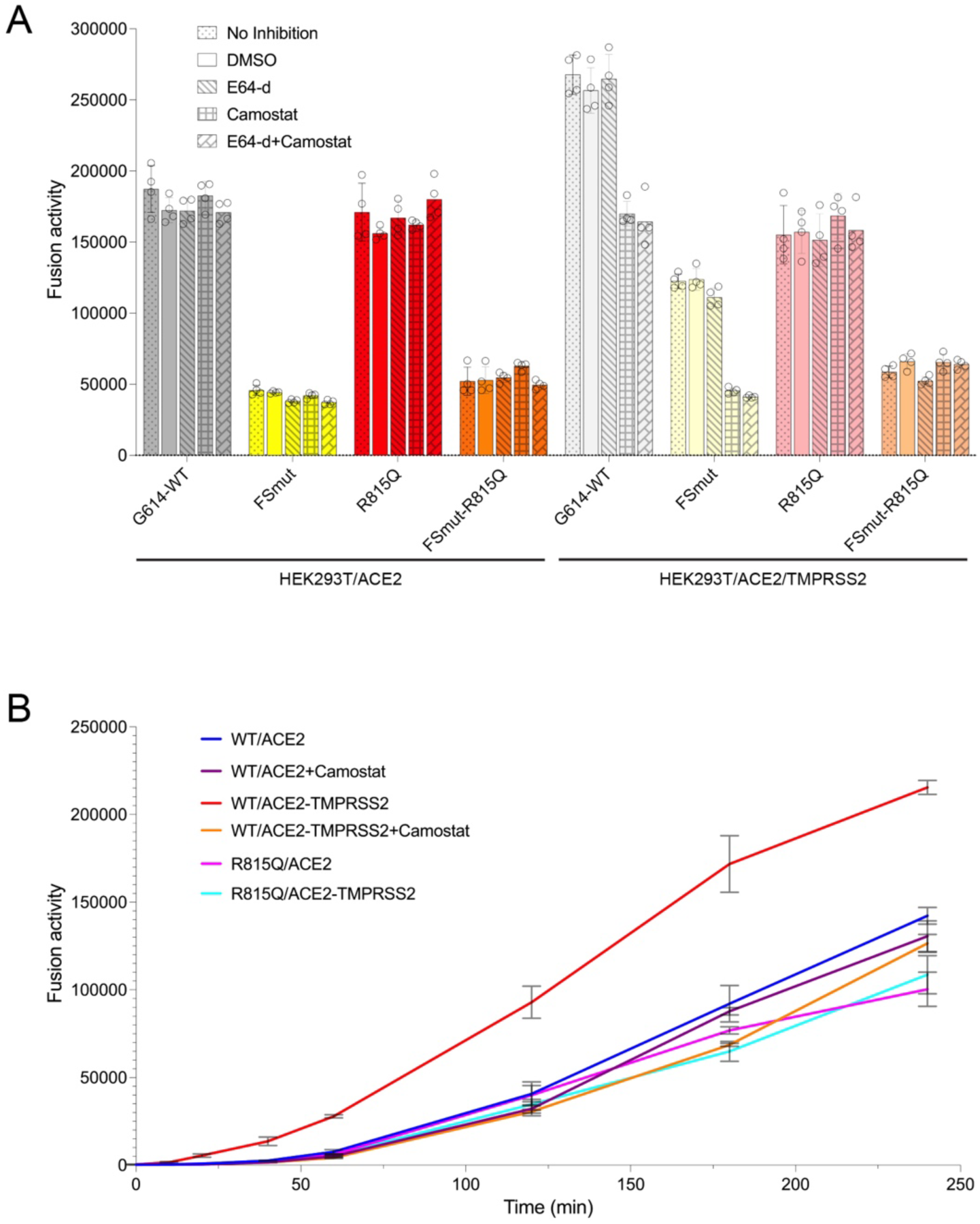
Functional roles of host proteases that cleave the S2’ site. (**A**) HEK293T cells transfected with the full-length S protein (G614) and selected mutants were tested for membrane fusion with the ACE2-expressing cells or ACE2- and TMPRSS2-expressing cells, in the presence of cathepsin L inhibitor E64-d or TMPRSS2 inhibitor camostat, or both. DMSO was used for dissolving E64-d and also tested as a control. The experiment was repeated at least three times with similar results. Error bars, mean ± s.d. (**B**) A time-course experiment for cell-cell fusion between HEK293T cells transfected with the full-length wildtype G614 S protein (WT) or the R815Q mutant and the ACE2-expressing cells or ACE2- and TMPRSS2-expressing cells. Camostat was also tested for blocking the effect caused by TMPRSS2. The experiment was repeated at least three times with similar results. Error bars, mean ± s.d.

We also performed a time-course experiment using the cell-cell fusion assay for a period of up to 4 hours, comparing the wildtype spike and R815Q mutant in the absence or presence of TMPRSS2 and camostat (Fig. 4B). The wildtype spike fused membranes much more rapidly in the presence of TMPRSS2 than it did in the absence of the protease in the target cells. The acceleration of fusion by TMPRSS2 was blocked by either comostat or by the mutation at the S2’ site.

## Discussion

The S2’ site cleavage has been considered essential to free the N-terminus of the segment immediately downstream (S^816-834^ for SARS-CoV-2), in order to allow its insertion into the target membrane and thus to function as a fusion peptide^31,33,39^. Our previous structure of the SARS-CoV-2 postfusion spike in membrane has demonstrated that it is the internal FP (residues 867-909), not the S^816-834^ segment, that spans the lipid bilayer^30^. In this study, using an S2 preparation with the S2’ site fully cleaved, we further show that three S^816-834^ segments of the trimer, with two of them fully visible and the other one partially disordered, are not buried in a hydrophobic environment, and thus are unlikely to serve as the fusion peptides. A recent study has reported an ACE2-induced early fusion-intermediate conformation of the SARS-CoV-2 spike, in which S2, not cleaved at the S2’ site, has partially refolded to eject the internal, cone-like FP cluster while S1 remains associated with S2 despite the furin cleavage (Fig. S9; ref^45^), further confirming our previous assignment of the FP for SARS-CoV-2 spike. These results suggest that the S2’ cleavage must fulfill some other purposes than releasing the N-terminus of the fusion peptide.

In our new structure, the three TM helices do not wrap around the trimeric FP cluster as they do in the uncleaved S2 postfusion trimer^30^. They instead run in parallel to each other, projecting laterally on one side of the trimer as part of a structure that appears to be stabilized by the S^816-834^ segments interacting with the TM-proximal part of HR2. This asymmetrical structure may represent a very late fusion intermediate conformation preceding the final, symmetrical postfusion conformation (Fig. S9). A recent cryo-ET study has identified several late fusion intermediates involving clusters of postfusion-like spikes after treatment with trypsin, presumably, the S2’ cleavage^46^. We cannot be certain that our new S2’-cleaved structure corresponds to one of these late intermediates because of the low resolution of the cryo-ET reconstructions. It is possible that in an asymmetrical structure, as shown in Fig. 2, with the FPs in the target membrane and the TM helices still in the viral membrane, the interactions between the FP-proximal S^816-834^ and the TM-proximal HR2 could bring the two lipid bilayers close together to induce formation of a hemifusion stalk^3,47^. Thus, the S^816-834^ segment, freed by the S2’ cleavage, might accelerate membrane fusion by promoting hemifusion and also by stabilizing this critical lipid intermediate. This role would be analogous to the capping structure near the membrane-interacting segments observed for influenza HA2^48^. Likewise, clustering of the S2’ trimers mediated by the FP-FP interactions could also help stabilize the hemifusion stalk and contribute to free energy required for the transition from hemifusion to fusion pore formation and dilation^3^.

Our mutational data demonstrate that R815, critical for the S2’ site cleavage at least by TMPRSS2 or trypsin, is not required for membrane fusion, since it can be substituted by many other residues as long as they do not disrupt the prefusion structure. The TMPRSS2-catalyzed cleavage increases the S-mediated membrane fusion activity and this enhancement can be equally abolished by either R815 mutation or TMPRSS2 inhibition, suggesting that the TMPRSS2 does not cleave at other Arg or Lys residues outside of the S2’ site. There is a substantial level of fusion activity that, while strictly dependent on both S and ACE2, is insensitive to the TMPRSS2 cleavage, indicating that there is little endogenous TMPRSS2 in HEK293T cells under the conditions that we used, and that other host protease(s) may have to cleave a site in the vicinity that is independent of R815. An intriguing study has mapped the cathepsin L cleavage sites in the SARS-CoV-2 spike to T259 and Y636, both in S1, and none in S2^49^, although cathepsin L has been generally assumed to be a surrogate of TMPRSS2 in endosomes to cleave the S2’ site. Further investigation is needed to clarify how these proteolytic events affects SARS-CoV-2 entry.

Finally, the S2’ site is targeted by several broadly neutralizing antibodies^50,51^, which could block either the cleavage or the HR2 folding back step to neutralize. The S2’ cleavage appears to break apart the epitope, which is also not very accessible in the ACE2-bound early fusion-intermediate conformation^45^. Thus, the uncleaved S2 postfusion structure with the S2’ site well-exposed should serve as an excellent immunogen to induce this type of broadly neutralizing antibody responses for development of pan-coronavirus vaccines.

## Methods

### Expression and purification of recombinant proteins

The expression constructs for the full-length spike (S) protein (residue 1-1273) of SARS-CoV-2 (G614) and monomeric soluble ACE2 protein were described previously^52,53^. The expression constructs for spike mutants were generated by standard PCR methods and verified by DNA sequencing of the entire coding region.

A stable cell line was generated for large-scale production of monomeric soluble ACE2 protein and purification of the ACE2 protein was carried out as previously described^53^. Briefly, the stably transfected cells expressing ACE2 were cultured in Expi293 expression medium (Thermo Fisher Scientific, Waltham, MA) containing 1% Pen Strep (Thermo Fisher Scientific) and 1.0 μg/ml puromycin (Thermo Fisher Scientific) to a density of ∼4-5×10^6^/ml, the cell supernatant was harvested by centrifugation at 3,000 ×g for 30 min and loaded onto a column packed with Ni Sepharose excel resin (Cytiva Life Sciences, Marlborough, MA). The column was washed with a buffer containing 20 mM Tris-HCl, pH 7.5, 300 mM NaCl and 10 mM imidazole. The ACE2 protein was eluted with a buffer containing 20 mM Tris-HCl, pH 7.5, 300 mM NaCl and 300 mM imidazole, and further purified by gel-filtration chromatography on a HiLoad 16/600 Superdex 200 pg column (GE Healthcare) in 25 mM Tris-HCl, pH 7.5 and 150 mM NaCl.

To produce the S2’ protein, we used a stable cell line generated using HEK293T cells for large-scale production of the full-length prefusion G614 S trimer as a starting material^54^. The stably transfected cells expressing S protein were grown in Expi293 expression medium containing 1% Pen Strep and 1.0 μg/ml puromycin to a density of ∼8×10^6^ cells/ml, and the cells were harvested by centrifugation at 2,000 ×g for 30 min. The cell pellets were washed with PBS and completely resuspended in PBS to the density of ∼8×10^6^ cells/ml. The resuspended cells were incubated with 0.5 μM soluble ACE2 protein and 27 μg/ml trypsin (Sigma-Aldrich, St. Louis, MO) at room temperature overnight. The cells were then lysed by adding 1% (w/v) n-dodecyl-β-D-maltopyranoside (DDM) (Anatrace, Inc. Maumee, OH), 1 mM phenylmethanesulfonyl fluoride (PMSF, Sigma-Aldrich, St. Louis, MO), EDTA-free complete protease inhibitor cocktail (Roche, Basel, Switzerland), and incubated at 4°C for 1 hour. After centrifugation at 27,000 ×g for 30 min, the supernatant was loaded onto a strep-tactin (IBA Lifesciences, Göttingen, Germany) column equilibrated with buffer A (100 mM Tris-HCl, pH 8.0, 150 mM NaCl, 1 mM EDTA) and 1% DDM. The column was washed with 50 column volumes of buffer A and 0.3% DDM, followed by additional washes with 50 column volumes of buffer A and 0.1% DDM, and with 50 column volumes of buffer A and 0.02% DDM. The S2’ protein was eluted by buffer A containing 0.02% DDM and 5 mM desthiobiotin (IBA Lifesciences), and further purified by gel-filtration chromatography on a Superose 6 10/300 column (GE Healthcare, Chicago, IL) in a buffer containing 25 mM Tris-HCl, pH 7.5, 150 mM NaCl, 0.02% DDM.

The S-specific monoclonal antibodies were purified as described previously^53,55^.

### Negative stain EM

To prepare grids for negative stain EM, 4 μl of freshly purified S2’ protein sample was adsorbed to a glow-discharged carbon-coated copper grid (Electron Microscopy Sciences, Hatfield, PA), washed with deionized water, and stained with freshly prepared 1.5% uranyl formate. Images were recorded at room temperature on a Phillips CM10 transmission electron microscope with a nominal magnification of 52,000×. Particles were auto-picked and 2D class averages were generated using RELION^56^ (4.0.1).

### Cryo-EM sample preparation and data collection

To prepare grids for cryo-EM, 4 μl of the S2’ protein sample at 4.5 mg/ml was applied to a 1.2/1.3 Quantifoil gold grid (Quantifoil Micro Tools GmbH), which had been glow discharged with a PELCO easiGlow^TM^ Glow Discharge Cleaning system (Ted Pella, Inc.) for 60 s at 15 mA. Grids were immediately plunge-frozen in liquid ethane using a Vitrobot Mark IV (ThermoFisher Scientific), and excess protein was blotted away using grade 595 filter paper (Ted Pella, Inc.) with a blotting time of 8 s and a blotting force of 5 at 4°C with 100% humidity. The grids were first screened for ice thickness and particle distribution. Selected grids were used to acquire images with a Titan Krios transmission electron microscope (ThermoFisher Scientific) operated at 300 keV and equipped with a Falcon 4i direct electron detector. Automated data collection was carried out using EPU version 3.8 (Thermo Fisher Scientific) at a nominal magnification of 165,000× in counting mode (calibrated pixel size, 0.736 Å) at an exposure rate of ∼8.13 electrons per pixel per second. Each movie adds a total accumulated electron exposure of ∼51.9 e-/Å^2^, fractionated in 59 frames. Data sets were acquired using a defocus range of 0.6-2.2 μm.

### Image processing and 3D reconstruction

Cryo-EM data processing was performed using cryoSPARC^37^ (v4.6.2) and RELION (5.0-beta3). Total 9969 movies were subjected to motion correction using patch mode, and contrast transfer function (CTF) estimated by patch mode as well in cryoSPARC. Based on a criterion of a CTF fit resolution exceeding 5Å, 9302 micrographs were selected, and subjected to template picking and Topaz picking^57^, yielding 902,903 particles. The selected particles were subsequently extracted with a box size of 256 pixels (1.73 A/pix sampling). After multiple rounds of 2D classification to eliminate junk particles, the remaining 603,296 particles were used to generate 5 *ab initio* classes, and heterogeneous refinement was performed to remove additional junk particles. Particles in the best class with clear structural features and detergent micelles were re-extracted without binning, giving 244,100 particles. A new *ab initio* model was then generated in cryoSPARC, and non-uniform refinement with C1 symmetry led to a 2.92Å map. The selected particles were then imported by Pyem tool^58^ into RELION for focused 3D classification with a mask encompassing the fusion peptide and transmembrane region. The best class from the focused 3D classification in RELION with 153,848 particles was re-imported into cryoSPARC to perform non-uniform refinement with C1 symmetry to produce a final consensus map of 3.08 Å resolution. Reference-based motion correction did not improve the resolution and was therefore not used for further processing. Instead, the best overall map was divided into two local maps: 1 and 2, using two masks generated for each part in cryoSPARC, to carry out particle subtraction and local refinement. For local map 1 (including the FP and TM region), initial local refinement yielded a 3.16Å map, which was subsequently improved to 3.04Å resolution after local CTF refinement. For local map 2, initial local refinement gave a 2.94 Å resolution map, which was subsequently improved to 2.82 Å resolution after local and global CTF refinement. The local maps were sharpened with deepEMhancer^59^ for model building. To confirm the presence of a class for the dimer of S2’ trimer in this data set, particles from the non-uniform refinement were aligned using the volume align tool in cryoSPARC to position two micelles in the center of the box to visualize the dimer interaction. A box size of 864 pixels was used to re-extract these particles, which were then binned by 2 to produce a box size of 432 pixels (1.472 Å/pix). 133,946 particles were subjected to non-uniform refinement and one round of 3D classification to give three classes. The class for the dimer of S2’ trimer was further refined by non-uniform refinement, producing a 4.19 Å map. In this map, the density for one trimer is substantially weaker than the other.

For a second independent data set, a total of 6,212 movies were imported and motion-corrected by patch mode in cryoSPARC. After CTF estimation, 5,917 micrographs were selected and subjected to template and Topaz picking, giving 191,567 particles. 185,919 of selected particles were extracted without binning, and subjected to multiple rounds of 2D classification to remove junk particles, leading to 73,177 particles. 5 *ab initio* models were generated and heterogeneous refinement performed to further clean up the particles. The best class was subjected to non-uniform refinement with C1 symmetry, producing a 4.22Å map, and further refined after reference-based motion correction in cryoSPARC to produce a 4.03Å map. Further classification led to two distinct classes. The class with a double-micelle feature, representing a dimer of trimer, was selected and refined with non-uniform refinement to produce a 4.28Å map. This map and its associated particles were centered around the micelle region using a volume alignment tool. These particles were re-extracted with a box size of 800 pixels, yielding 14,159 particles, and the final non-uniform refinement produced a 6.80Å map. In this map, the difference in density for two trimers is smaller than that in the map from the first data set.

All resolutions were reported from the gold-standard Fourier shell correlation (FSC) using the 0.143 criterion. Local resolution was also determined using cryoSPARC.

### Model building

For model building, our previously published model for the uncleaved S2 trimer in membrane (PDB: 8FDW)^30^ was used as a template. The new model was fitted in ChimeraX^60^ and manually adjusted in Coot^61^. ModelAngelo^62^ was used to help model the regions that are different from our previous S2 structure. The complete S2’ model was refined in Phenix^63^ using real-space refinement with geometric restraints, and validated in both Phenix and MolProbity^64^.

### Western blot

Full-length S protein or S2’ protein samples were resolved in 4-15% Mini-Protean TGX gel (Bio-Rad) and transferred onto PVDF membranes (Millipore, Billerica, MA) by an Iblot2 (Invitrogen by Thermo Fisher Scientific). Membranes were blocked with 5% skimmed milk in 1× PBST for 1 hour. To detect the expression and cleavage at S1/S2 furin site of full-length S protein, the SARS-CoV-2 (2019-nCoV) Spike RBD Antibody (Sino Biological Inc., Beijing, China) at the concentration of 1 μg/ml were incubated for another one hour at room temperature. Alkaline phosphatase conjugated anti-Rabbit IgG (1:5000) (Sigma-Aldrich, St. Louis, MO) was used as a secondary antibody. To detect the S2’ protein with intact C-terminal twin Strep tag, the Strep-Tactin AP conjugate (1:4000) (IBA Lifesciences) was used as the primary antibody. To detect the cleavage at S2’ site, a human monoclonal antibody CC40.8^65^, which targets the conserved stem helix epitope of S2, at the concentration of 1 μg/ml were incubated for another one hour at room temperature. Alkaline phosphatase conjugated anti-human IgG (1:5000) (Jackson ImmunoResearch, West Grove, PA) was used as a secondary antibody. To detect nucleocapsid protein, 2 µg/mL anti-SARS-CoV-2 Nucleocapsid Mouse Monoclonal IgG (R&D Systems, Minneapolis, MN) was used as the primary antibody. Alkaline phosphatase conjugated goat-anti-mouse IgG (1:5000) (Jackson ImmunoResearch, West Grove, PA) was used as a secondary antibody. Proteins were visualized using Western Blue® Stabilized Substrate for Alkaline Phosphatase (Promega, Madison, WI).

### Cell-cell fusion assay

The cell-cell fusion assay, based on the α-complementation of *E. coli* β-galactosidase, was carried out to quantify the fusion activity mediated by SARS-CoV-2 S protein, as described previously^27^. Briefly, the full-length G614 S or its mutants (10 μg) and the α fragment of *E. coli* β-galactosidase construct (10 μg), or the full-length ACE2 construct (5 μg) (without or with 5 μg of the full-length TMPRSS2 construct) together with the ω fragment of *E. coli* β-galactosidase construct (10 μg), were transfected in HEK293T cells using polyethylenimine (PEI) (80 μg). After incubation at 37°C for 5 hrs, the medium was aspirated and replaced with complete DMEM (1% Pen Strep, 1% GlutaMax and 10% FBS), followed by incubation at 37°C for additional 19 hrs. The cells were detached using PBS and resuspended in complete DMEM. 50 μl S-expressing cells (1.0×10^6^ cells/ml) were mixed with 50 μl ACE2-expressing cells (1.0×10^6^ cells/ml) to allow cell-cell fusion to proceed at 37°C for 4 hrs. For assays with protease inhibitors, 25 μM E-64d (Sigma-Aldrich, St. Louis, MO) or 100 μM camostat mesylate (Sigma-Aldrich, St. Louis, MO) were used. Cell-cell fusion activity was quantified using a chemiluminescent assay system, Gal-Screen (Applied Biosystems, Foster City, CA), following the standard protocol recommended by the manufacturer. The substrate was added to the mixture of the cells and allowed to react for 90 min in dark at room temperature. The luminescence signal was recorded with a Synergy Neo plate reader (Biotek, Winooski, VT).

### SARS-CoV-2 VLP infectivity assay

SARS-CoV-2 virus-like particles (VLPs) were produced following a protocol reported previously^42^, with minor modifications. Briefly, HEK293T cells (in 10 cm plates) were transfected with SARS-CoV-2 spike, membrane, nucleocapsid and envelope expression constructs, as well as the construct PS9 expressing a viral RNA segment for packaging into VLPs and a luciferase reporter (Luc-PS9/Plasmid#177942, Addgene; ref^42^). At 22 hours post-transfection, the cell culture medium was replaced, and at 48 hours post-transfection, the cell supernatant was harvested, filtered with 0.45 micron filter, and then incubated with Retro-X™ Concentrator (Takara Bio USA, Inc., San Jose, CA) overnight at 4°C. The supernatant was centrifuged at 1500 xg for 45 min to harvest precipitated VLPs, which were then resuspended in 100 μl PBS as the VLP stock. To measure infecitivty of VLPs, 50 μl of cell suspension containing 3×10^4^ target cells (HEK293T cells transfected with ACE2 alone or ACE2 and TMPRSS2) and 50 μl of medium containing 2.5 μl of each SARS-CoV-2 VLP stock were loaded in each well of an opaque white 96-well plate. The plate was incubated at 37°C for 26 hours, allowing VLP infection and luciferase expression. The cell supernatant was removed from each well, and the cells rinsed with PBS and subsequently lysed in 20 μl passive lysis buffer (Promega) for 15 minutes at room temperature with gentle rocking. 50 μl of reconstituted luciferase assay buffer was added and luminescence was measured immediately after mixing using a BioTek Synergy Neo microplate reader (BioTek Instruments, Winooski, VT).

### Flow cytometry

Expi293F cells were grown in Expi293 expression medium. Cell surface display DNA constructs for the SARS-CoV-2 G614 or its mutants or S2 together with a plasmid expressing blue fluorescent protein (BFP) were transiently transfected into Expi293F cells using ExpiFectamine 293 reagent (Thermo Fisher Scientific) according to the manufacturer’s instruction. 2 days post-transfection, the cells were stained with primary antibodies at a concentration of 5 μg/ml. An Alexa Fluor 647-conjugated donkey anti-human IgG Fc F(ab′)2 fragment (Jackson ImmunoResearch) was used as the secondary antibody at a concentration of 5 μg/ml. Cells were run through an Intellicyt iQue Screener Plus flow cytometer, and those gated for the positive BFP expression were analyzed for antibody binding.

## Acknowledgments

We thank S. Harrison for critical reading of the manuscript, and the SBGrid team for computing support. We acknowledge support for COVID-19 related structural biology research at Harvard from the Nancy Lurie Marks Family Foundation and the Massachusetts Consortium on Pathogen Readiness (MassCPR). This work was supported by COVID-19 Awards from MassCPR (to B.C.), and NIH grant AI127193 (to B.C. and S. Harrison).

## Author Contribution

B.C. and W.S. conceived the project. W.S. produced the S2’ protein, prepared cryo grids and performed EM data collection with contributions from R.M.W.. GM.J. and W.S. processed the cryo-EM data, built and refined the atomic model. S.W. and J.L. created the stable cell lines. H.Z. performed the flow cytometry experiment. W.S. created the mutant constructs, and carried out the cell-cell fusion assay. G.K. produced SARS-CoV-2 VLPs and performed the VLP infectivity assay. J.A., H.P. and S.R.V. contributed to plasmid preparation, cell culture and antibody production. All authors analyzed the data. B.C., W.S. and GM.J. wrote the manuscript with input from all other authors.

## Competing Interests

All authors declare no competing interests.

## Data Availability

The atomic structure coordinates and EM map are deposited in the EMDataBank under the accession number: PDB ID 9NXY, EMD-49912, EMD-49917, EMD-49918, and EMD-49921. All other related data generated during and/or analyzed during the current study, such as raw cryo-EM images, are available from the corresponding authors on reasonable request. Source data are also available.

## Notes

### Competing Interest Statement

The authors have declared no competing interest.

