## Supplementary materials for "Effect of the S2’ site cleavage on SARS-CoV-2 spike"

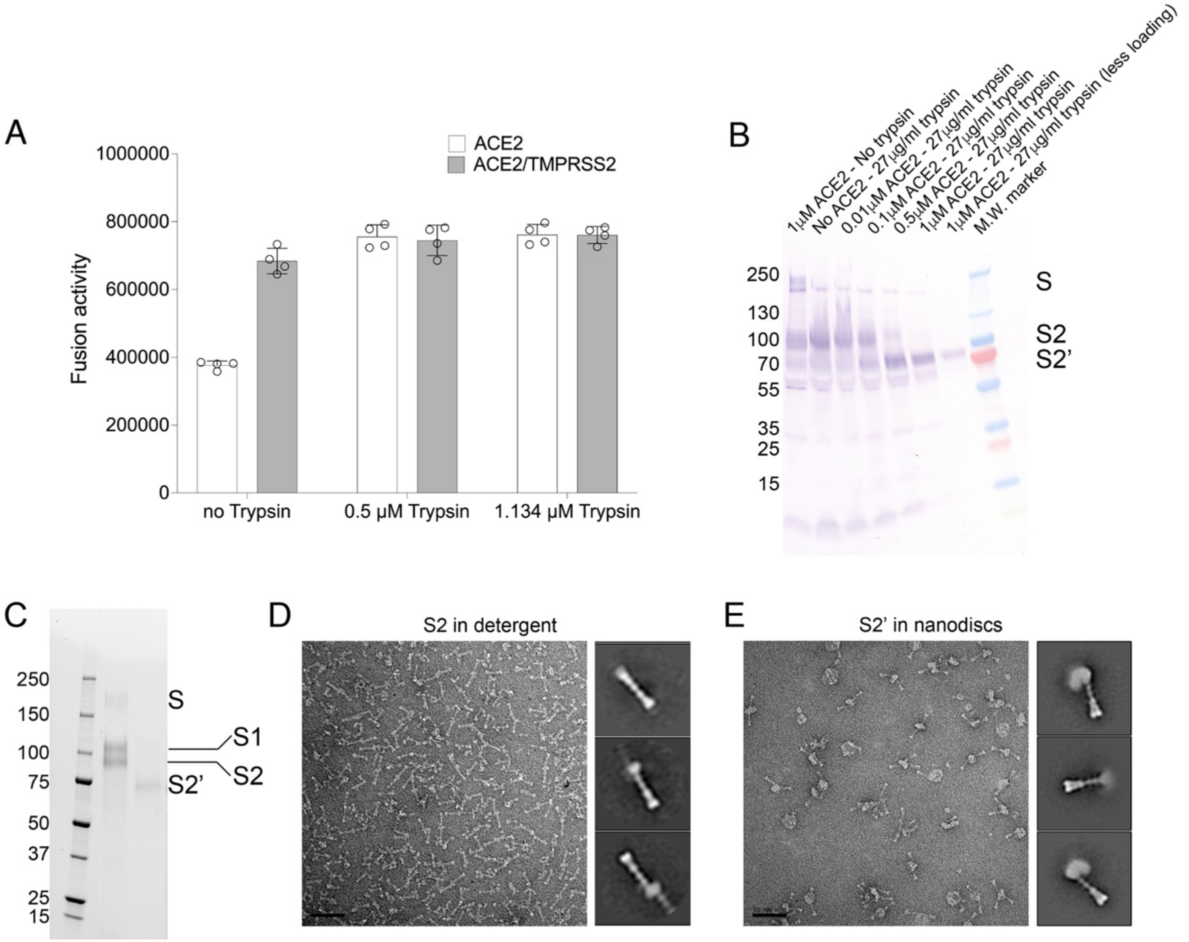

**Figure S1. Production of the S2' fragment of SARS-CoV-2 spike.** (**A**) Both TMPRSS2 and trypsin enhance S-mediated membrane fusion. HEK293T cells transfected with the full-length S protein (G614) were tested for membrane fusion with the ACE2-expressing cells or ACE2- and TMPRSS2-expressing cells in the presence of trypsin. The experiment was repeated at least three times with similar results. (**B**) Cleavage of the S protein by trypsin in the presence of ACE2. S-expressing cells were treated with soluble ACE2 and trypsin at various concentrations as indicated and their lysates were analyzed by western blot using anti-strep tag antibody. (**C**) SDS-PAGE analysis showing the purified S2' fragment migrated faster than the uncleaved S2, as expected. (**D**) Negative stain EM of the uncleaved S2 trimers in detergent showing clusters of trimers. Representative image and 2D averages are shown, and the box size of 2D averages is ~460Å. (**E**) Negative stain EM of the S2' trimers reconstituted in nanodiscs showing clusters of trimers. Representative image and 2D averages are shown, and the box size of 2D averages is ~460Å.

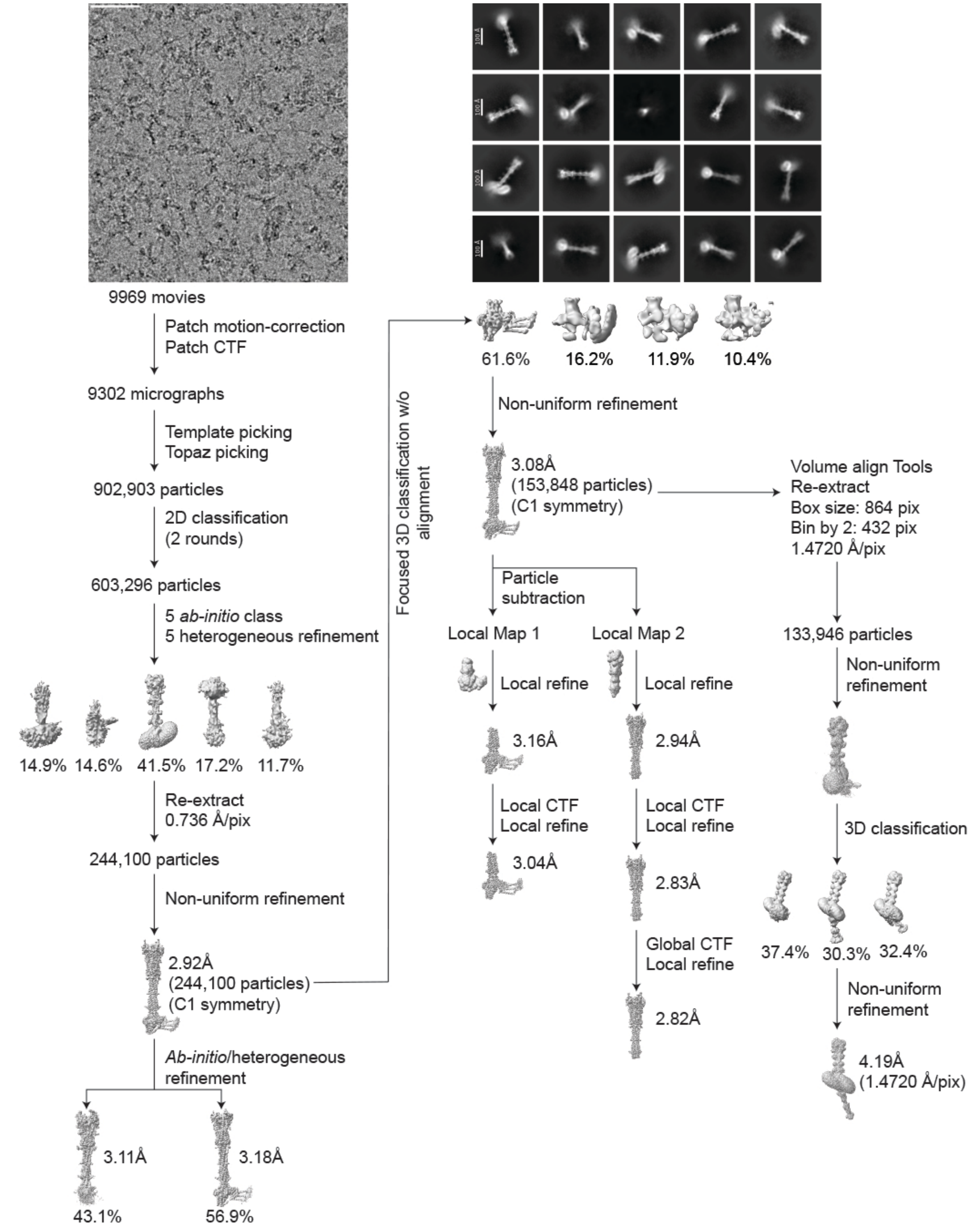

**Figure S2. Cryo-EM analysis of the S2' trimer.** Top, representative micrograph out of 9,969 similar micrographs**,** and 2D averages of the cryo-EM particle images of the postfusion S2' trimer in detergent. Bottom, data processing workflow for structure determination.

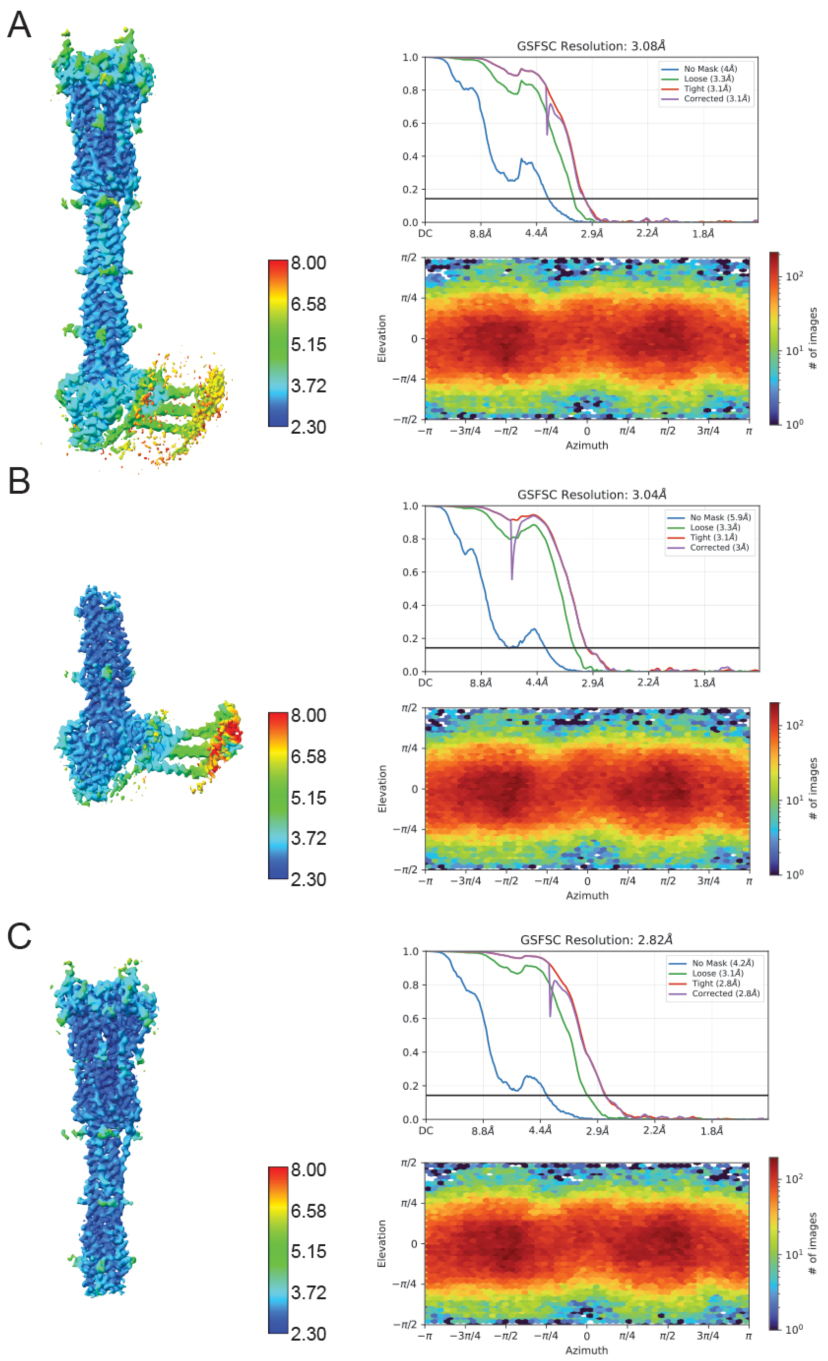

**Figure S3. Analysis of the S2' trimer structures by cryoSPARC.** (**A**)-(**C**) 3D reconstructions of the S2' trimer from the overall refinement and local refinement (top and bottom) are colored according to local resolution estimated by cryoSPARC with the FSC=0.5 criterion. Gold standard FSC curves of the three refined 3D reconstructions of the S2' trimer and the corresponding cryoSPARC output for particle distribution of each reconstruction.

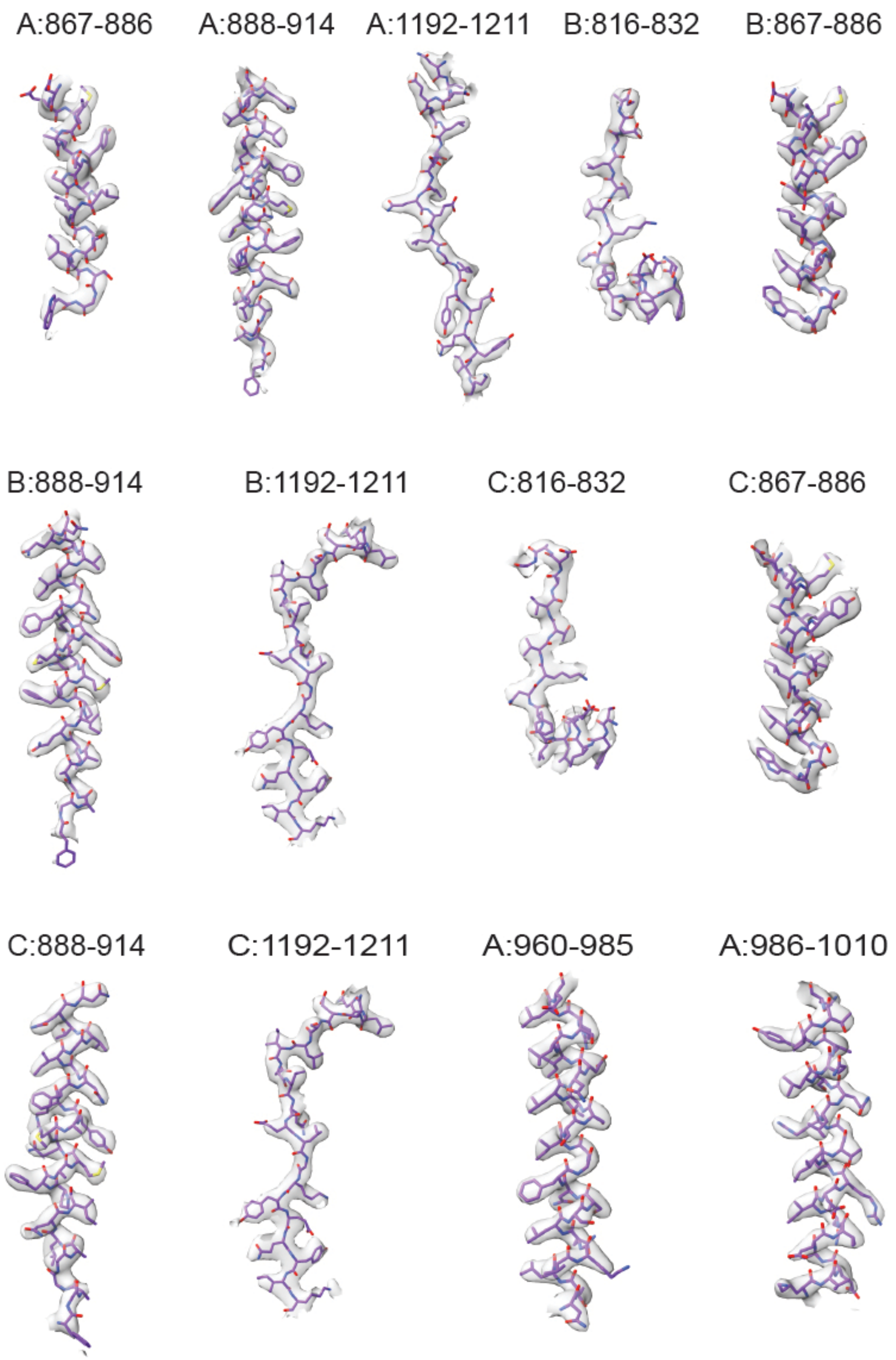

**Figure S4. Additional analysis of the S2' trimer structure.** Representative density in gray surface representation from the EM map of the S2' trimer. The model range is indicated by chain name and residue numbers. For example, A:867-886 represents chain A: residues 867-886.

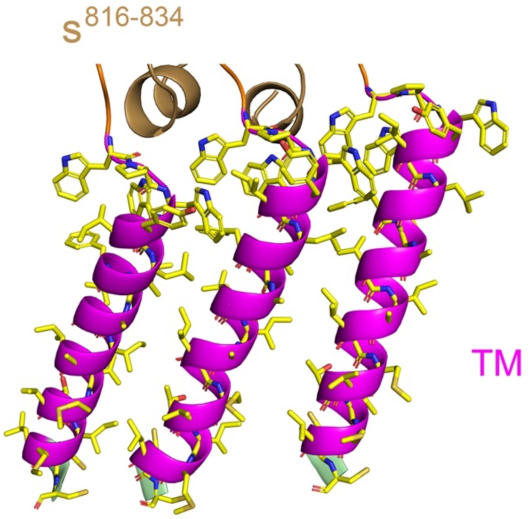

**Figure S5. TM helices in the S2’ trimer.** A close-up view of the three TM helices in the S2’ trimer in detergent micelle.  All the residues are shown in stick model.

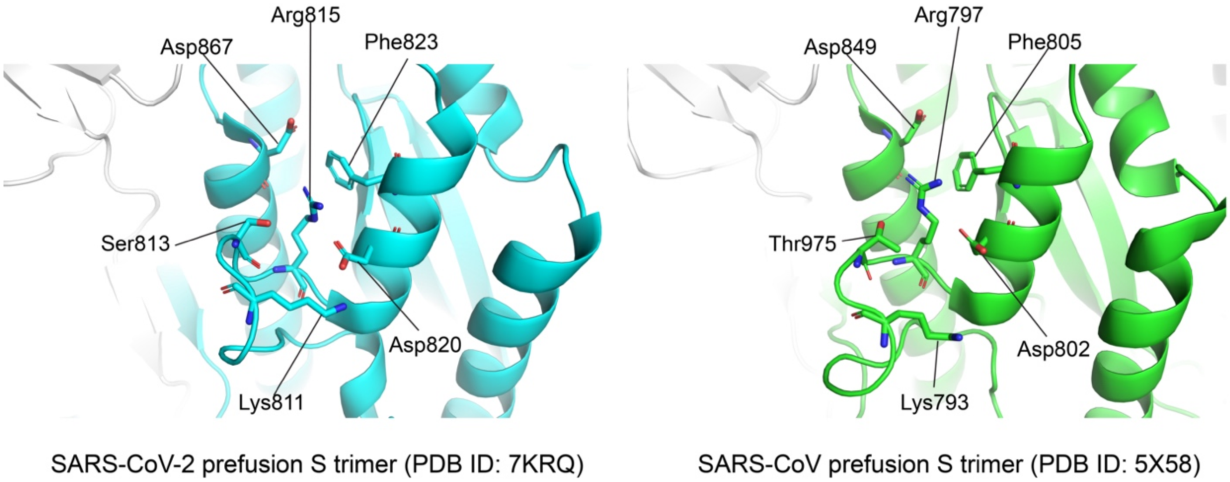

**Figure S6. Local structures near the S2' cleavage site in SARS-CoV-2 and SARS-CoV prefusion spike trimers.** The prefusion trimer structures of SARS-CoV-2 (PDB ID:7KRQ[52]) and SARS-CoV (PDB ID:5X58[66]) are shown in ribbon diagram. The residues near the S2' cleavage site (Arg815 in SARS-CoV-2 and Arg797 in SARS-CoV) are indicated.

A

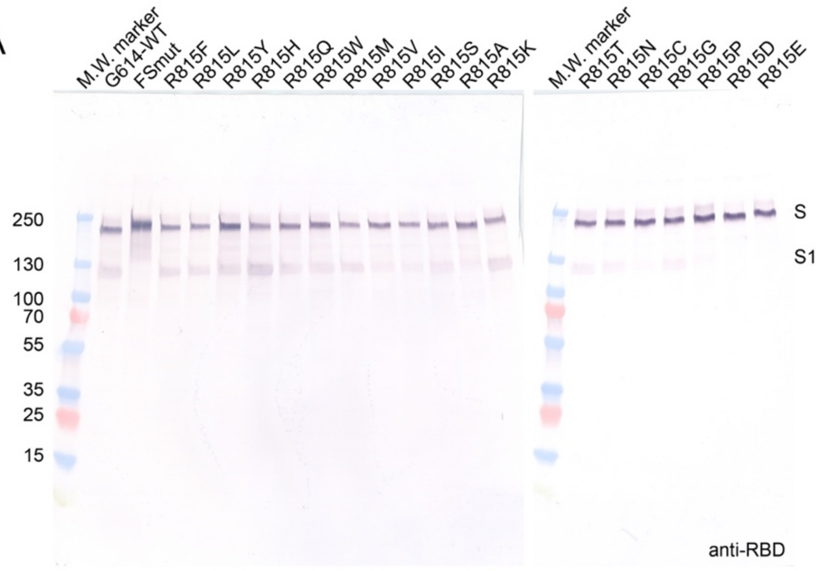

B

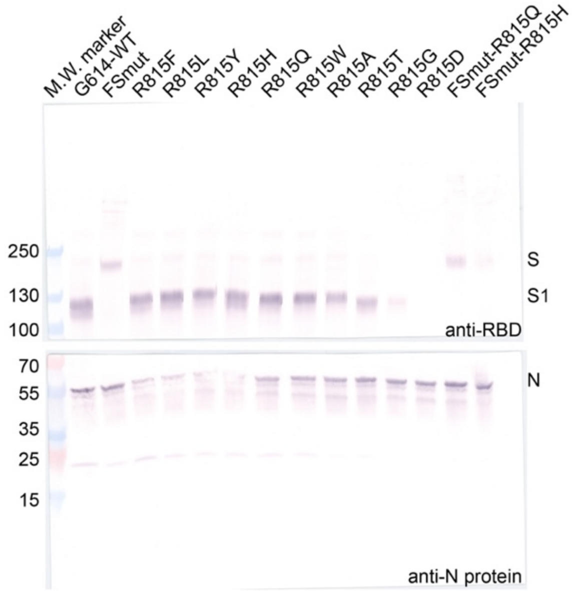

**Figure S7. Expression of the mutant spike proteins and their incorporation into SARS-CoV-2 VLPs.** (**A**) Expression and processing of the full-length G614 S construct and its mutants in HEK293 cells. S protein samples prepared from HEK293 cells transiently transfected with 10 μg of the full-length S expression plasmids were detected by anti-RBD polyclonal antibodies. Bands for the uncleaved S and S1 fragment are indicated. The experiment was repeated twice with similar results. (**B**) VLP samples using G614 S construct and selected mutants were detected by anti-RBD polyclonal antibodies and an anti-N (nucleocapsid) protein. Bands for the uncleaved S, S1 fragment and N protein are indicated. The experiment was repeated twice with similar results.

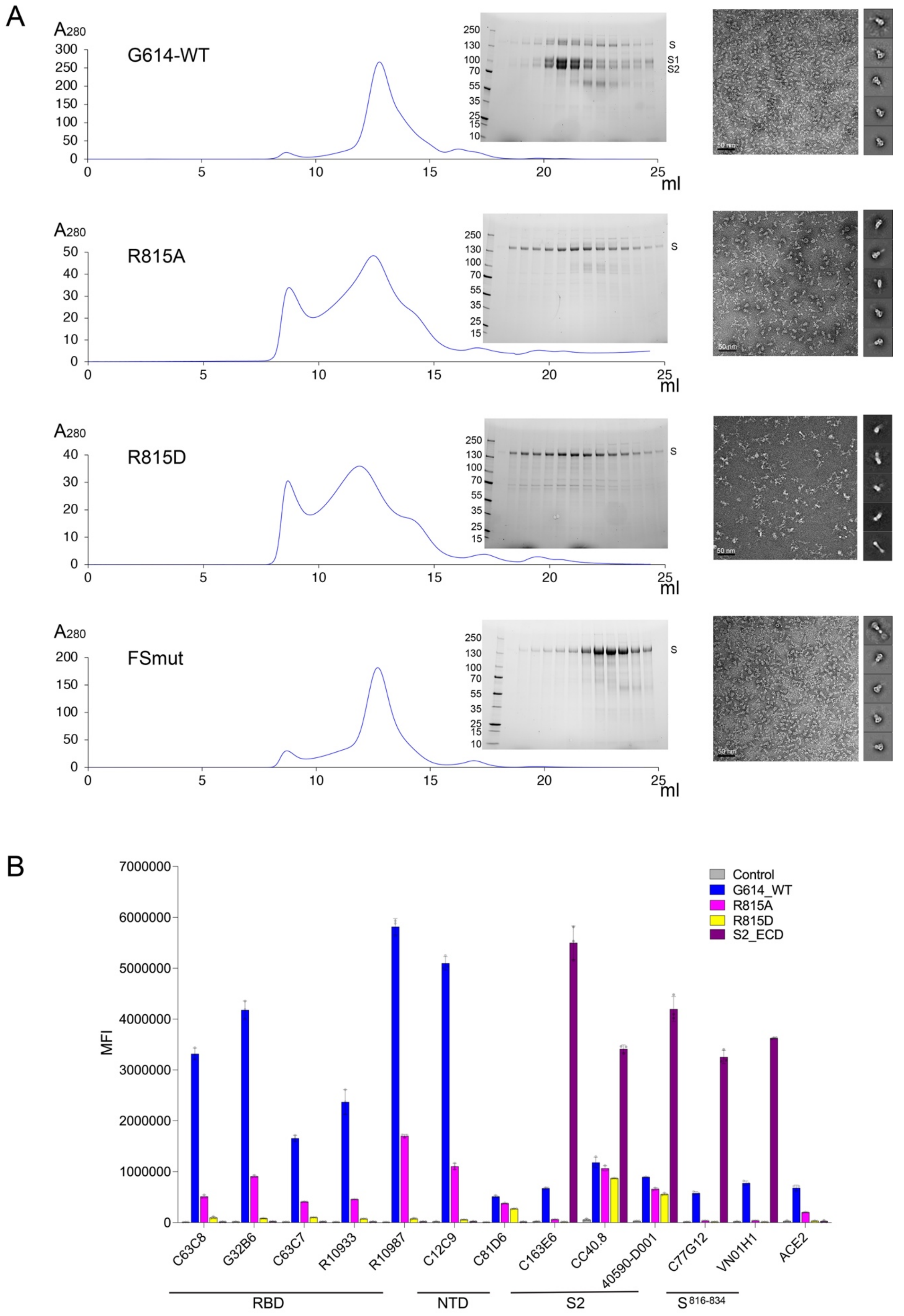

A
A280
G614-WT
R815A
R815D
FSmut
ml
S
S1
S2
250
130
100
70
55
35
25
15
10
50 nm
B
MFI
Control
G614_WT
R815A
R815D
S2_ECD
C63C8
G32B6
C63C7
R10933
R10987
C12C9
C81D6
C163E6
CC40.8
40590-D001
C77G12
VN01H1
ACE2
RBD
NTD
S2
S 816-834

**Figure S8. Characterization of mutant spike proteins.** (**A**) The purified full-length SARS-CoV-2 S proteins with mutations as indicated were resolved by gel-filtration chromatography on a Superose 6 column. Peak fractions were analyzed by SDS-PAGE. Representative 2D averages by negative stain EM of the peak fractions are also shown. The box size of 2D averages is ~460Å. G614-WT, R815A and FSmut showed predominantly the prefusion trimers, while R815D showed the postfusion trimers without any recognizable prefusion trimers. (**B**) Antibody binding to the full-length G614 S protein and its mutants R815A and R815D, as well as an S2 construct (S2_ECD) expressed on the cell surfaces analyzed by flow cytometry. The antibodies and their targets are indicated. A designed ACE2-based fusion inhibitor $ACE2_{615}$-foldon-T27W was used for detecting receptor binding[53]. MFI, mean fluorescent intensity. The error bars represent standard errors of mean from measurements using three independently transfected cell samples. The flow cytometry assays were repeated three times with essentially identical results.

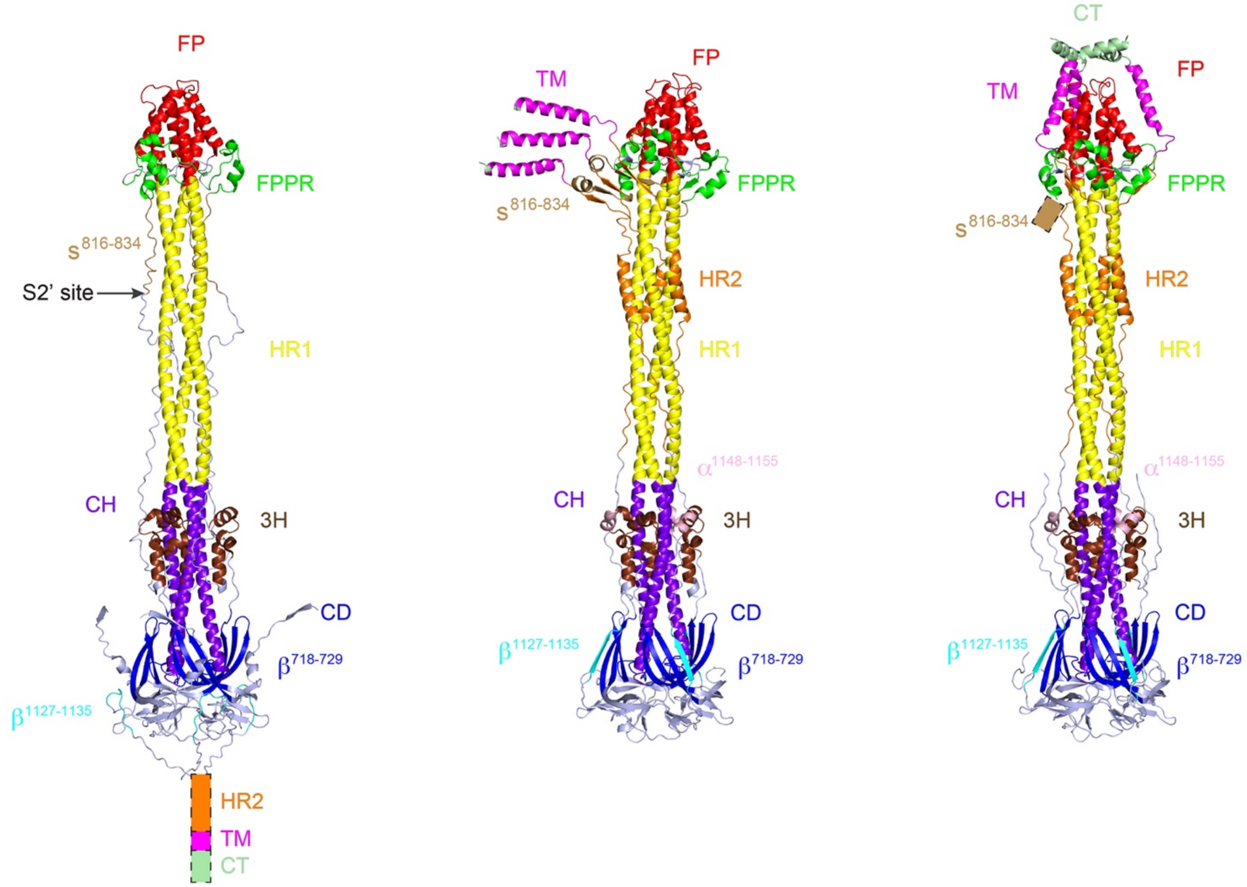

**Figure S9. Structural comparison of the early fusion intermediate S2 trimer, S2' trimer and uncleaved postfusion S2 trimer.** Structures of the early fusion intermediate S2 trimer (PDB ID: 8Z7P[45]), S2' trimer (this study) and uncleaved postfusion S2 trimer (PDB ID: 8FDW[30]) are shown in ribbon diagram. Various structural components include $\beta^{718\text{-}729}$, 3H, $s^{816\text{-}834}$, FPPR, FP, HR1, CH, CD, $\beta^{1127\text{-}1135}$, $\alpha^{1148\text{-}1155}$, HR2, and CT.

**Table S1. Cryo-EM data collection, refinement and validation statistics**

| | SARS-CoV-2 S2' protein (EMD-49912) (PDB ID 9NXY) | SARS-CoV-2 S2'-localmap1 (EMD-49917) | SARS-CoV-2 S2'-localmap2 (EMD-49918) |
|---|---|---|---|
| **Data collection and processing** | | | |
| Magnification | 165,000 | 165,000 | 165,000 |
| Voltage (kV) | 300 | 300 | 300 |
| Electron exposure (e–/Å$^2$) | 51.929 | 51.929 | 51.929 |
| Defocus range (μm) | 0.6-2.2 | 0.6-2.2 | 0.6-2.2 |
| Pixel size (Å) | 0.736 | 0.736 | 0.736 |
| Symmetry imposed | C1 | C1 | C1 |
| Initial particle images (no.) | 902,903 | | |
| Final particle images (no.) | 153,848 | 153,848 | 153,848 |
| Map resolution (Å) | 3.08 | 3.04 | 2.82 |
| FSC threshold | 0.143 | 0.143 | 0.143 |
| Final map sampling (Å/pix) | 0.736 | 0.736 | 0.736 |
| **Refinement** | | | |
| Initial model used (PDB code) | 8FDW | | |
| Model resolution (Å) | 3.3 | | |
| FSC threshold | 0.5 | | |
| Model resolution range (Å) | | | |
| Map sharpening *B* factor (Å$^2$) | | | |
| Model composition | | | |
| Non-hydrogen atoms | 11992 | | |
| Protein residues | 1463 | | |
| Ligands | NAG:39,MAN:15 | | |
| *B* factors (Å$^2$) | | | |
| Protein (min/max/mean) | 29.46/227.90/88.24 | | |
| Ligand (min/max/mean) | 96.08/240.05/166.69 | | |
| R.m.s. deviations | | | |
| Bond lengths (Å) | 0.005 | | |
| Bond angles (°) | 0.495 | | |
| Validation | | | |
| MolProbity score | 1.70 | | |
| Clashscore | 5.97 | | |
| Poor rotamers (%) | 1.92 | | |
| Ramachandran plot | | | |
| Favored (%) | 97.11 | | |
| Allowed (%) | 2.89 | | |
| Disallowed (%) | 0.00 | | |
